# Reusing excess staple oligonucleotides for economical production of DNA origami

**DOI:** 10.1101/2024.08.27.609958

**Authors:** Giorgia Isinelli, Christopher M. Wintersinger, Pascal Lill, Olivia J. Young, Jie Deng, William M. Shih, Yang C. Zeng

**Author notes:** These authors contributed equally to this work (Giorgia Isinelli, Christopher M. Wintersinger, Yang C. Zeng).

## Abstract

DNA origami has enabled the development of responsive drug-delivery vehicles with precision features that were previously not attainable in bionanotechnology. To reduce the costs of creating therapeutics-scale amounts of DNA origami that need to bear costly modifications with high occupancy, we reused the excess staple oligonucleotides that are leftover from the folding process to fold additional origami. We determined that a DNA origami can be successfully folded with up to 80% cost savings by cyclic recovery and reuse of excess staple strands. We found evidence that higher quality staple strands are preferentially incorporated into origami, consistent with past reports, and therefore are preferentially depleted from the free-strand pool. The folding of DNA origami with staple strands that were reused up to eleven times was indistinguishable by our panel of assays versus a control folded with new strands, so long as the reused oligonucleotides were replenished each cycle with a small excess of fresh strands. We also observed a high degree of incorporation of guests on the DNA origami. By recovering, reusing, and replenishing excess staple oligonucleotides, it is possible to significantly lessen production costs to create well-formed origami, which is useful to allow more therapeutic designs to be tested.

## INTRODUCTION

DNA origami is a proven and robust method for forward design and synthesis of two- and three-dimensional nanoscale shapes for responsive drug-delivery vehicles^1^. In the approach, an excess of complementary “staple” DNA oligonucleotides^2^ anneal to a long ssDNA “scaffold,” with strategically-placed crossovers to fold the scaffold into the designed shape. One promising application of origami is as vaccines^3–5^ and for drug delivery, where the terminal ends of staple oligonucleotides are used to precisely position therapeutic cargoes at nanometer spacing intervals^3,6,7^. Such programmability has enabled drug delivery vehicles that release their payload when triggered by *in vivo* stimuli and vaccines with tunable immune responses^8–10^. However, the therapeutic use of origami has been impeded by a number of challenges, including two key ones that need to be considered together, highlighted below.

Firstly, the cost of manufacture of large amounts of origami bearing staple strands with expensive modifications presents an impediment for its application as therapeutics^11^. This expense is amplified by the requirement for providing large excesses of staple strands over scaffold to ensure the highest incorporation efficiency. Milligram scales of product as needed to prototype potential therapeutics are most readily made using staple oligonucleotides from chemical-synthesis vendors, which typically represent the majority of the material cost of DNA-origami fabrication (see Supplementary Text 1.1 for further discussion, from Table S2 to Table S5).

Secondly, reliable and repeatable results with therapeutic origami require folding conditions that favor high incorporation rates of the designed features. It is notable that the robustness of origami folding, in terms of preferential incorporation of higher-quality staple strands, allows production of well-formed origami even with unpurified commercially purchased staple strands. For instance, a typical commercial synthesis with solid support phosphoramidite chemistry has a yield upwards of ∼0.99^(*n*-1)^ for an *n* length strand, such that about one-third of the product has critical errors (e.g. missing an important 5′ end cargo feature) for a typical 42mer staple strand. Folding is resilient to impurities of the staple-strand pool because of cooperative processes favoring the addition of full-length strands. Nonetheless, there is still only ∼50–95% incorporation rate of any particular 3′ feature using typical reaction conditions, in the case of folding with a 10-fold excess of staple strands^12^. Larger staple excesses or purification of the input oligonucleotides can further improve the incorporation of desired 3′ and 5′ features, but this would require greater amounts of starting materials and therefore would further increase the cost of production for applications such as therapeutics.

To address this unmet need for lower cost origami therapeutics, here we have explored reusing the excess staple strands leftover from folding to synthesize more origami in the context of milligram-scale production suitable for animal studies (Fig. 1A). This can yield substantial savings on material cost for staple strands, with greater cost reductions as excess strands are reused repeatedly to fold additional rounds of origami. With a ten-fold excess of staple strands in the folding reaction, the unit cost of oligonucleotides for the origami could be nearly halved with only a single cycle of staple reuse, versus using only freshly purchased staple strands. Moreover, the total oligonucleotide costs could be lessened by ∼80% with ten rounds of reuse or when larger excesses of staple strands are used (Fig. S1, Supplementary Text 1.2). We initially observed that the staple strands left over from folding could be efficiently recovered by ethanol precipitation from a supernatant containing the remaining staple strands after recovery of the origami product by polyethylene glycol (PEG) precipitation (Fig. S2, Method 4, Method 5). Encouraged by this technique, we tested whether the reuse of staple strands would allow the formation of DNA origami designed for a therapeutic application with satisfactory incorporation of 3′ and 5′ features on the product.

**Figure 1.**
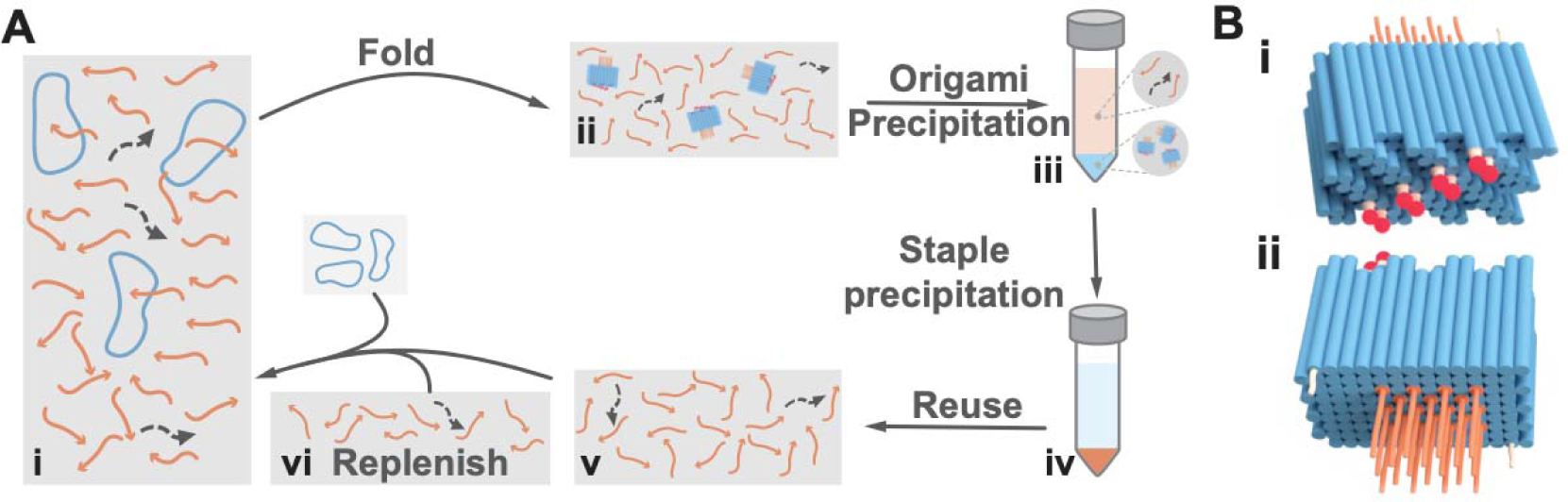
Proof-of-concept therapeutic DNA origami SQB and an approach for folding it by reusing excess staple strands. Ai-ii, Folding of the SQB from the circular scaffold DNA 50nM (blue) and excess unpurified staples strands 250 nM core staples, 500 nM handles and 1000 nM for functional staples (with full-length strands in orange versus truncated impurity strands in dark grey). **Aiii**, PEG precipitation is used to separate the DNA origami (in the pellet) from the unused staple strands (in the supernatant). **Aiv**, Ethanol precipitation followed by centrifugation is used to extract the unused staple strands (into the pellet) from the PEG purification buffer. **Av**, The folding process favors incorporation of full-length strands into the origami over their truncated counterparts, reducing the average purity of the remaining strand pool. **Avi,** The recovered strands may be reused to fold more DNA origami, if they are replenished with an equivalent of strands to what were consumed in the preceding folding experiment. We posit that replenishing the reused strands maintains full-length strands at sufficient concentration, such that staple strands may be reused multiple cycles to fold high-quality origami. **Bi–ii**, Renderings of the front and back cargo-holding faces of the DNA origami square block with CpG and fluorophores conjugated.

One pitfall we foresaw was that the relative purity of the staple strands might be progressively lessened with reuse if folding preferentially incorporates full-length staple strands^2^, or if long incubations as elevated temperatures lead to accumulation of significant depurination event^13^. We hypothesized that replenishment of the recovered staple pool with fresh staple strands, to replace each equivalent of staple strands which were removed in the prior folding cycle, would sufficiently restore the average strand quality to an extent that enables folding of additional origami of similar quality to those from earlier cycles (Fig. 1Avi, Fig. S3, see Supplementary Text 1.1 for discussion about the DNA origami costs).

## MATERIAL AND METHODS

### Method 1, SQB DNA origami folding

Unpurified dehydrated staple oligonucleotides were purchased from Integrated DNA Technologies (IDT) at either 10 or 100 nmol scales and rehydrated in water so that the concentration of each strand was ∼0.2 or ∼0.5 mM. Oligos for a particular origami were combined using volumes to create 2× staple strand stocks, with the concentrations of each strand specified in Table S1. The p8634 scaffold strand was produced from M13 phage replication in JM109 *Escherichia coli*. Large scale (∼40 mL) folding mixtures were prepared in ten parts total as follows: five parts 2× staple strand stock, one part folding buffer (50 mM Tris pH 8.0, 10 mM EDTA, 120 mM MgCl_2_), with the remaining four parts to add the appropriate scaffold at 50 nM final concentration, Cy5 labeling strand at the concentration noted in Table S1, and water. The larger reaction mixture was split into smaller volumes in 0.2 mL PCR tubes and incubated on a thermocycler as follows: 80°C/15 minutes in a single step; 50–40°C in 100 10.8 min steps less 0.1°C/step; 4°C thereafter until collection of the sample for further processing and PEG purification^6^.

### Method 2, Barrel DNA origami folding

Unpurified dehydrated staple oligonucleotides were purchased from Integrated DNA Technologies (IDT) at either 10 or 100 nmol scales and rehydrated in water so that the concentration of each strand was ∼0.2 or ∼0.5 mM. Oligos for a particular origami were combined using volumes to create 2× staple strand stocks, with the concentrations of each strand specified in Table S6. The p7308 scaffold strand was produced from M13 phage replication in JM109 *Escherichia coli*. DNA origami folding mixtures were prepared in ten parts total as follows: five parts of the 2× staple stock, one part of folding buffer (50 mM Tris pH 8.0, 10 mM EDTA, 120 mM MgCl_2_), with the remaining four parts to add the appropriate scaffold at 50 nM final concentration, Cy5 labeling strand at the concentration noted in Table S6, and water. The larger reaction mixture was split into smaller volumes in 0.2 mL PCR tubes to be incubated on a thermocycler as follows: 80°C for 15 minutes, decrease the temperature to 60–25°C in 350 3.08 min steps less 0.1°C/step; 4°C thereafter until collection of the sample for further processing and PEG purification^14^.

### Method 3, Nanocube folding and purification

The design for a 10 nm × 10 nm × 10 nm nanocube was adapted from as previously published^15^. Unpurified dehydrated nanocube oligonucleotides were purchased from Integrated DNA Technologies (IDT) at a 10-nmol scale, rehydrated in water at ∼100 μM each, and mixed with equal volumes per strand. Out of the 28 total nanocube strands, one strand was selected and appended with a 4T linker and 16-nt handle to its 3′ end. The nanocube was prepared with ∼1 μM of each strand (except for the handle tagged strand at ∼2 μM), 40LmM MgCl_2_, and folded with a 42-hour temperature gradient: 80°C/10 min in a single step; 65–37°C in 290 8.69 min steps less 0.1°C/step; 16°C thereafter until collection of the sample.

The purification method used was not the one indicated in the original article because it provided a solution with a yield that was too low for the experiments. The nanocubes have been precipitated with 20% PEG precipitation in a 1:1 ratio volume, MgCl_2_ balanced. The resulting pellet was resuspended keeping into consideration that for the experiment the concentration had to be high enough. Then we performed agarose-gel-based extraction. With this combined method, a concentrated solution of pure product was obtained.

### Method 4, PEG precipitation to purify origami from excess staple strands

The following protocol was adapted from as previously published^16^. For the SQB, one volume of a PEG purification buffer (5 mM Tris pH 8.0, 1 mM EDTA, 15% w/v PEG-8000, 510 mM NaCl, 8 mM MgCl_2_) was added to an equal volume of the raw folded SQB in a 50 mL conical tube. For the barrel, the raw folding reaction was diluted ten-fold (in 5 mM Tris, 1 mM EDTA, 12 mM MgCl_2_). Next, one volume of diluted barrel folding reaction was added to ten volumes of a PEG purification buffer (5 mM Tris, 1 mM EDTA, 10% w/v PEG-8000, 510 mM NaCl, ∼12 mM MgCl_2_) in a 15 mL conical tube. The sample was mixed thoroughly by aspirating and dispensing it using a pipette, incubated for 30 minutes in a dark setting, and then placed on a centrifuge at 16k x g for 25 minutes at room temperature. The majority of the supernatant was separated from the pellet containing the SQB by either decanting or removing it with a pipette. Next, the sample was centrifuged for an additional two minutes at 16k x g so that the remaining supernatant could be gently extracted using a pipette. We note that the pellets appeared blue, because of the Cy5 fluorophore attached to the SQB. A volume of buffer (5 mM Tris pH 8.0, 1 mM EDTA, 10 or 12 mM MgCl_2_ for the SQB and barrel respectively) was added to dilute the origami at the approximate desired concentration and incubate for five minutes at 37°C. The pellet was detached from the wall of the tube by flicking it, at which point the sample was placed on a shaker at 600 rpm, 37°C for 25 minutes. The sample was cooled to room temperature and then the origami concentration was determined by measuring the amount of DNA on a Nanodrop 2000c spectrophotometer.

### Method 5, Recovery of excess staple oligonucleotides from PEG supernatant

The leftover staple supernatant from PEG purification was reserved from the last step of Method 4. We recorded the total volume of supernatant and used a Nanodrop 2000c spectrophotometer to quantify the starting amount of ssDNA. In four parts total, one part of staple supernatant was mixed thoroughly with three parts of ethanol (100% or 200-proof). Next, one-tenth of the staple supernatant volume of 3 M sodium acetate was added. The solution was cooled in a -80°C freezer for one hour and then put in a chilled centrifuge at 21k × g for 30 minutes. The majority of ethanol supernatant was decanted from the tube into another one. Then the tube was briefly centrifuged so the last traces of supernatant could be removed with a pipette. We proceeded to wash the pellet with a tenth of the total previous volume with cold ethanol (75% v/v). Then the tube was centrifuged for 5 minutes at the same speed and temperature as the previous step. The ethanol was decanted and removed with a pipette as described above, and the wash step was repeated one more time. The pellet was left to air dry in a clean place, with the total drying time selected depending on the relative size of the pellet (i.e. 15 min for small pellets, vs. overnight for the largest pellets). Sterile water was added to the dried pellet in a volume to dilute the staple strands to the approximate desired concentration. The final concentration of the strands was determined on a Nanodrop 2000c spectrophotometer, such that the recovery yield could be determined with respect to the starting measurement.

We note that this yield was necessary for the amounts of replenishing strands in Method 6. Also, salt contamination of the recovered staple strands was problematic for the successful folding of the SQB. In cases where the 260/230 absorbance ratio was much larger than ∼2.2, we concluded that contaminating salts of the oligonucleotides was too high. In these occurrences, ethanol precipitation of the sample was repeated as above until a satisfactory 260/230 ratio was attained.

### Method 6, Typical approach for replenishment of the recovered staple strands

This approach is appropriate if certain staple strands have different relative excesses versus other staple strands with respect to the scaffold, in a given folding mixture. Recovered staple strands from Method 5 were replenished with an equivalent amount of fresh strands to replace strands extracted by the origami in the prior folding or that were lost in the recovery procedure. The following considerations were made in replenishing the staple strands: One, each type of SQB staple is added in differing stoichiometric excesses with respect to the scaffold (Table S1); Two, the scaffold during the previous folding removed one equivalent of staple strands from the total strand excess; And three, there was a marginal loss of staple strands during recovery in Method 5. As such, the volume of each staple type mixture to add for replenishment was determined as below. The *original_volume* refers to the volume added initially to create the 2× staple strand stock during the initial folding with fresh strands in Method 1, and the *recovery_yield* was determined in Method 5.

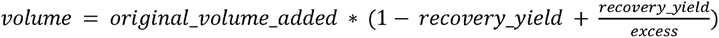

The replenishing strands were added to the recovered staple mixture in the volumes as computed above, with the mixture diluted with water to attain the 2× staple strand stock volume that was added during reaction setup for the previous folding. We proceeded to set up the folding reaction using the replenished staple mixture as described in Method 1. Additionally, we added one-fifth of the amount of Cy5 labeling strand initially used in Method 1.

### Method 7, Alternative approach for replenishment of the staple strands

This approach is appropriate if all the staple strands have the same relative excesses with respect to the scaffold in a given folding mixture. Initially, we used a Nanodrop 2000c spectrophotometer to determine the initial ssDNA concentration of the 2× staple stock mixture used in Method 1. The excess staple strands as recovered in Method 5 were rehydrated in a minimal volume of sterile water, and were diluted in more water to match the concentration of the initial 2× staple stock mixture. The final volume of the recovered diluted strands was recorded, with an additional fresh 2× staple stock mixture added to restore the volume of staple stock used during the prior folding cycle. Finally, we prepared the folding reaction with the replenished staple mixture as described in Method 1.

### Method 8, Negative-stain transmission electron microscopy (TEM)

Origami samples were diluted to a final concentration of 4 nM in 1× TEF buffer (5 mM Tris pH 8.0, 1 mM EDTA, 10–12 mM MgCl_2_). For the SQBs samples, we used a 1×TEF buffer with 10 mM MgCl_2_ to dilute. For the nanocube samples alone and the SQBs conjugated with nanocubes, we used a 1×TEF buffer with 10mM MgCl_2_. For the barrel samples we used 1×TEF buffer with 12mM MgCl_2_ to dilute and for the conjugated product Barrel-SQB we used 1xTEF buffer with 11mM MgCl_2_. TEM grids (Electron Microscopy Sciences FCF400-CU-50, or alternatively grids that we carbon-coated in ourselves) were negatively glow discharged at 15 mA for 25 s in a PELCO easiGlow. The diluted sample (4 µL) was applied to the glow discharged grid, incubated for 2 min, and wicked off completely into Whatman paper (Fisher Scientific, 09-874-16B). Immediately after, 10 µL of 2% aqueous filtered uranyl formate, was applied for 1-2 s, immediately wicked off. The procedure was repeated and this time the uranyl formate was left for 40–45 s and then dried against Whatman paper leaving a very thin layer on the grid. Grids were left to dry in a dust-free area. All imaging was performed at 80kV on JEOL JEM 1400 plus microscope and captured with AMT Image Capture Engine Software Version 7.0.0.255. Micrographs have been collected at a nominal magnification of 30k and pixel size 0.4nm/px for SQB samples conjugated with Barrels and for SQB conjugated with Nanocubes nomina magnification is 40k with 0.3nm/px as pixel size. Conjugation analysis of cargoes to the SQB was performed using the EMAN 2.11 software package^17^. Respective particles have been selected and extracted using the e2boxer tool. Particular analysis has been performed by displaying the selected particles with e2display.

All images presented in this work were imported into FIJI ImageJ (v1.53c)^18^, corrected for background noise using a pseudo-flat-field image, and then contrast and brightness adjusted for clarity in publication.

### Method 9, Agarose gel electrophoresis and gel yield calculations

Gel characterization of DNA origami samples was performed using the Thermo Scientific™ Owl™ EasyCast™ B2 electrophoresis system. UltraPure agarose (Life Technologies, 16500500) was melted in 0.5× TBE (45 mM Tris, 45 mM boric acid, 0.78 mM EDTA, 11 mM MgCl_2_) to a concentration of 2.0 % (w/v). The molten agarose was cooled and SYBR Safe was added (10 μL/160 mL of molten agarose). The samples were prepared in 10 μL volumes as follows: 5 μL of 6× AGLB (5 mM Tris, 1 mM EDTA, 30% w/v glycerol, 0.025% w/v xylene cyanol) loading dye, 250 fmoles of the sample, and sterile water to reach the final volume. Control sample lanes were generally ∼0.5 μg of GeneRuler DNA Ladder 1 kb 250–10,000 bp ladder (ThermoFisher SM0311). The mixed samples were loaded onto the gel and separated for ∼2 hours at 70 V at room temperature. Gel images were captured on a GE Typhoon FLA 9000 fluorescent imager using the SYBR Safe parameters as given in the Typhoon control software, with analysis of the images performed with FIJI ImageJ (v1.53c)^18^. Background subtraction with a rolling ball radius of 30–60 pixels was performed on linear TIFF images. The GelAnalyzer plugin in ImageJ and wand tool were used to integrate total pixel intensities from lanes of interest. DNA origami yields were determined by taking the ratio of the intensity of the band of interest with respect to all the species with a molecular weight larger than the excess staple strands.

### Method 10, Agarose-gel-based extraction

Five parts of the folded DNA nanocube were combined with one part of 6× AGLB (5 mM Tris, 1 mM EDTA, 30% w/v glycerol, 0.025% w/v xylene cyanol) loading dye. The samples were loaded and run on agarose gels as described in Method 9. The bands were observed on a UV transilluminator and cut from the gel using a razor blade. The band slices were placed in a 15 mL tube and then thoroughly crushed using a large pestle (BioMasher V, Funakoski Co.). The tubes were inverted on a centrifuge, spun at 1.5× g for one minute, at which point the crushed gel could be transferred from the tube lid to a DNA spin column (Freeze ‘N’ Squeeze, Bio-Rad) with tweezers. The gel slice was further crushed using a small disposable pestle against the walls of the DNA spin column tube and then centrifuged at 7k × g for 5 min at room temperature. The flow-through solution containing the DNA structure was transferred to another collection tube. Next, the gel slices were again disturbed with the pestle, and the spin column again centrifuged as above, with the remaining flow combined into the other collection tube.

### Method 11, CpG loading efficiency

The DNA origami was digested with DNase I so that the amount of nuclease-resistant CpG strand could be measured to determine the relative amount of the cargo loaded. Using Thermo Fisher DNase I, RNase-free kit (1U/μL, EN0521), the reaction mixture was prepared in a 0.2 mL PCR tube by mixing the purified SQB (2 μg), 10× DNase I reaction buffer (1 μL), DNase I enzyme (1.5 μL), and water to attain a final 10 μL volume. Control samples with only the CpG staple oligonucleotides with and without DNase I was also prepared. The reactions were incubated at 37°C for 30 minutes to fully digest the sample. One part of the sample was mixed with one part of formamide loading buffer (2× solution of 95% Formamide, 18 mM EDTA, and 0.025% SDS, Xylene Cyanol, and Bromophenol Blue) which was then heated to 94°C for 2 minutes and cooled to 4°C until analysis on a gel. The samples were loaded onto a 15% denaturing polyacrylamide gel, which was prepared using the SequaGel UreaGel System (National Diagnostics, EC-833) and plastic 1.0 mm mini-gel cassettes (Invitrogen NovexTM, NC2010). Samples were migrated into the gel at 250 V for 45 minutes in 0.5× TBE (45 mM Tris, 45 mM boric acid, 0.78 mM EDTA) buffer. The gel was stained for 15 minutes in SYBR Gold (i.e. 3 μL of stain added to 15 mL of MilliQ water) on a horizontal shaker table in a dark room. Gel images were captured on a GE Typhoon FLA 9000 fluorescent imager using the SYBR Gold parameters as given in the Typhoon control software, with densitometry of the images performed with FIJI ImageJ (v1.53c)^22^ to determine relative CpG loading. (Fig. S5)

### Method 12, Conjugation of nanocubes and barrels to the SQB

The purified SQB was mixed with either purified nanocube (at ten-fold stoichiometric excess to ∼50 nM SQB) or origami barrel (at two-fold stoichiometric excess to ∼25 nM SQB) in 1× TEF buffer (5 mM Tris pH 8.0, 1 mM EDTA, ∼12 or 10 mM MgCl_2_ for the nanocube and barrel respectively). The samples were heated for one hour at 37°C on a shaking incubator. In the case of samples conjugated to the nanocube, excess nanocubes were removed from the sample using PEG purification (as described for the SQB in Method 4, Fig. S7). Subsequently, images of the sample were captured by TEM so that the number of bound cargoes on single particles could be counted manually.

## RESULTS AND DISCUSSION

We studied the folding of a DNA origami square block (i.e. SQB) used as a proof-of-concept vaccine nanoparticle in order to determine the viability of staple-strand reuse^6^. The dimensions of the SQB are ∼35 nm × ∼27 nm × ∼22.5 nm, with its square-lattice arrangement presenting 126 modifiable 3′ and 5′ helical ends, which can be appended with therapeutic cargoes. The flat face in Fig. 1Bi displays 18 CpG immunostimulant strands with 3.5 nM spacing, recognized as the optimal spacing for Toll-Like Receptor 9 (TLR9) interaction^6,19–22^. We note that we employ CpG segments bearing a non-natural phosphorothioated backbone so as to protect these sequences from nuclease digestion for *in vivo* environments. Conversely, the jagged face in Fig. 1Bii has single-stranded DNA (ssDNA) handle strands to bind Cy5 fluorophores and other cargoes for downstream experiments.

We first tested folding of the SQB through five cycles of reuse of the staple strands, in this case using a two-fold molar excess of the latter over scaffold strand in each cycle sans any replenishment with fresh staple oligonucleotides. The volume of each folding reaction with reused staple strands was lessened to maintain the relative two-fold excess of staple strands. As shown on the agarose gel in Fig. 2Ai, the SQB has similar mobility after a single cycle of reuse as compared to the origami when folded only with fresh staple strands. However, there is a drastic slowdown in the gel mobility of the structure by the second cycle of reuse, which becomes even more extreme by the third, fourth, and fifth rounds. Gel densitometry indicated that the relative yield of the SQB monomer was diminished from ∼60% to ∼20% between the first and second reuse cycles, and finally with negligible yield after additional reuse cycles (white circular data points on Fig. 2B, see Method 9 for details about yield calculations). Using negative stain transmission electron microscopy (TEM), we concluded that well-formed SQB particles could not be observed by the third cycle of staple reuse without replenishment (Fig. 2Ci–ii). These observations suggest that the origami folding process preferentially selects full-length strands from a folding solution containing truncated staple oligonucleotides; a similar argument has been made earlier to rationalize the observation that a large excess of synthetic staple strands is needed for high-quality folding of DNA origami more generally^2^. We concluded that the staple pool was gradually enriched with undesirable flawed strands with continued cycles of reuse, to the extent that it could no longer allow satisfactory folding of the SQB.

**Figure 2.**
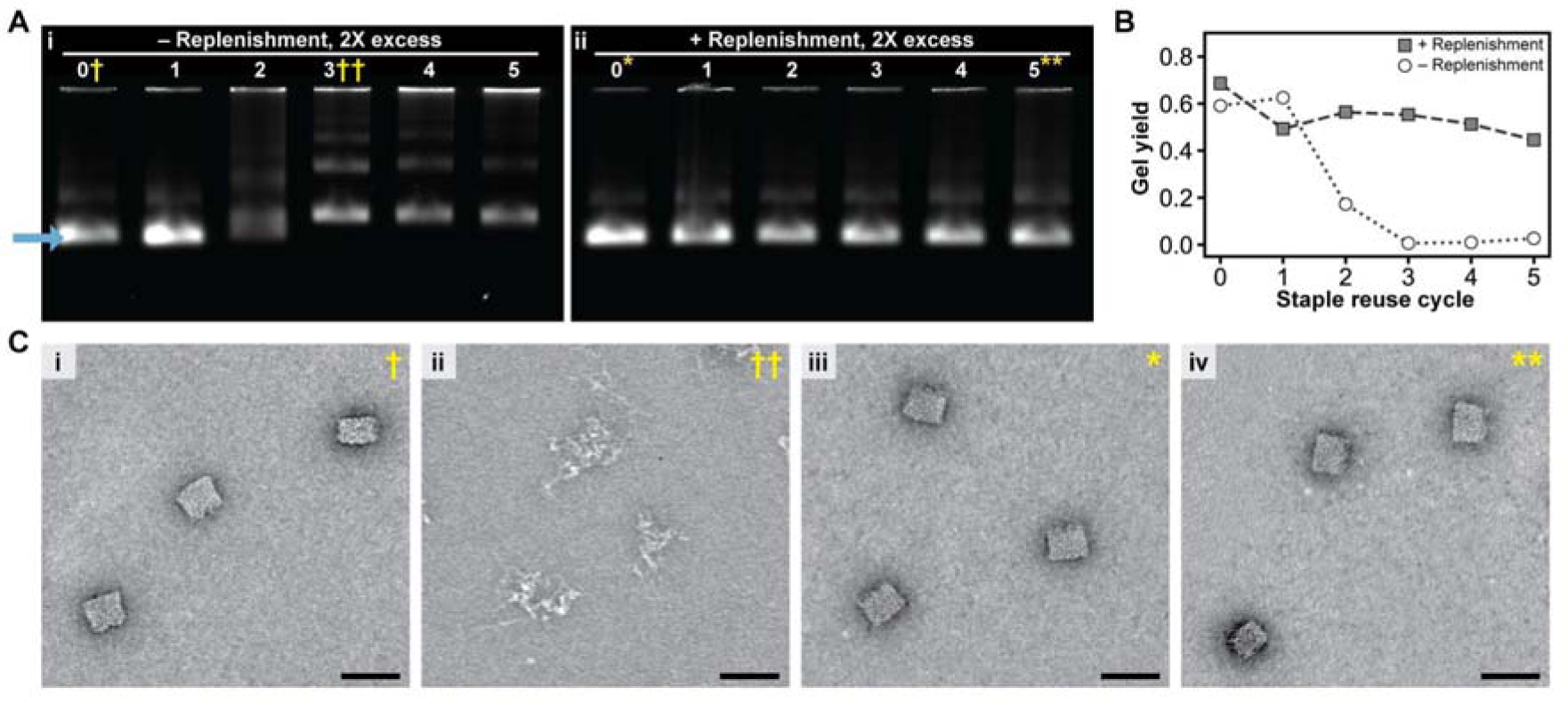
Replenishment of staple strands in excess with fresh strands enables successful folding of the DNA origami therapeutic with multiple cycles of reuse. **A**, Agarose gels of the purified origami product that were folded with a 2× excess of staple strands. There is a drastic decrease in the mobility of the structure after two or more cycles of reuse when the staple-strand pools are not replenished with fresh strands (**i**), versus similar mobility with reuse with replenishment (**ii**). **B**, Yields of the desired monomer structure, with the yield determined from densitometry of the SQB band (i.e. the blue arrow) with respect to the overall well. **C**, Representative negative-stain TEM micrographs showing single folded SQB particles of the samples noted on the gel in *A*. Scale bars are 50 nm.

To see whether staple strands with consecutive cycles of reuse could be used to properly fold the SQB, we repeated the aforementioned experiment with replenishment of these strands. After each folding using a two-fold excess of staple strands, the excess strands were collected and combined with one equivalent of fresh strands to replace those that were incorporated into the origami in the previous folding round. On an agarose gel, we observed similar mobility of the SQB regardless of whether it was folded with fresh staple strands versus if the strands were reused and replenished up to five times, with ∼60% yield of the desired structure (Fig. 2Aii, gray square data points on Fig. 2B). While there did seem to be a gradual and slight diminishing of the yield with more reuse of the staple strands (e.g. ∼45% by the fifth cycle), observation of single SQB particles by TEM indicated qualitatively similar folding between the initial and last cycles of reuse (Fig. 2Ciii–iv). Taken together, these observations suggest that replenishment of the recovered staple-strands pool with fresh strands can sufficiently restore these pools over multiple cycles of reuse so that they can satisfactorily enable the folding of the SQB.

Having shown the viability of folding origami with staple strands that were in excess, we wanted to see whether satisfactory folding could be attained over a greater number of cycles of reuse. We supposed that larger excesses of staple strands would be more robust to depletion of full-length strands over continued reuse because there would be a larger total starting pool of oligonucleotides. We note that we initially focused on a mere two-fold excess of staple strands over scaffold for easier testing of our hypothesis about the eventual need for replenishment (i.e. to make this more evident at an earlier cycle of staple-strand reuse). Given that larger excesses of staple strands increase the likelihood that 3′ and 5′ features are exhibited^12^, and the importance of high- incorporation rates of such features for making the effect of an origami therapeutic reliable and repeatable, we folded the SQB using the following molar excesses as determined by their therapeutic importance: (1) immunostimulant CpG staple strands at twenty-fold excess; (2) ssDNA handle strands for Cy5 fluorophores and other potential cargos (eg. antigens) at ten-fold excess; and (3) core strands for the structural integrity of the SQB at five-fold excess (Table S1). Indeed, reuse and replenishment of staple strands could sustain satisfactory folding of the SQB with ∼80% yield across eleven cycles of reuse, as shown by the agarose gel and densitometry of the gel in Fig. 3A–B. Single SQB particles were qualitatively similar when folded using fresh staples, versus with staples reused four, eight, or eleven cycles, as shown in the TEM micrographs in Fig. 3i–iv. We also note that yield of SQB was noticeably diminished after more than six cycles of reuse with no replenishment using the standard staple excesses noted above, when the total volume of reaction was not adjusted to maintain the standard excess of staple strands with respect to the scaffold (Fig. S4).

**Figure 3.**
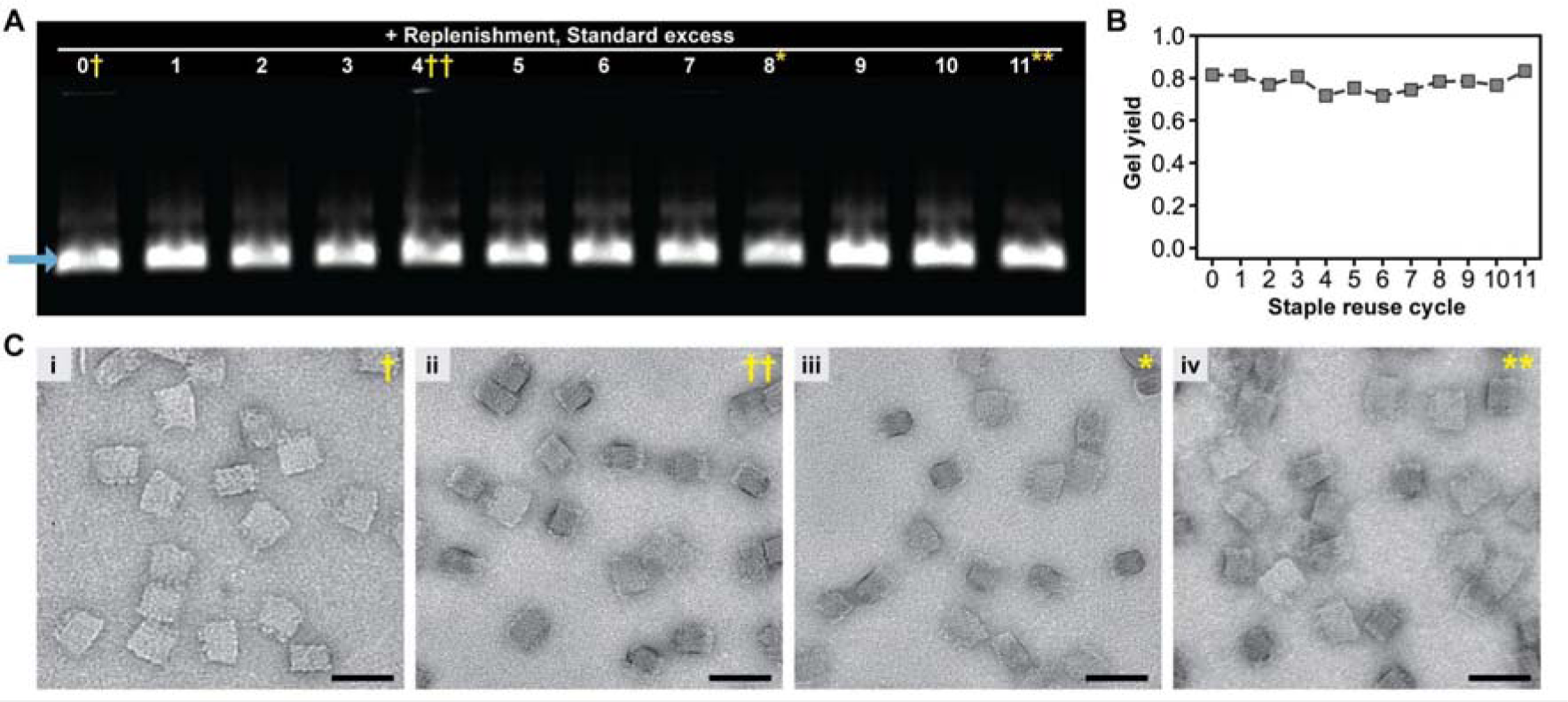
**A–B** Agarose gel showing that folding yield is sustained with replenishment of reused staple strands when there is at least a 5× excess of staple strands, with the yield determined from densitometry of the SQB band (i.e. the blue arrow) with respect to the overall well. **C**, Representative negative-stain TEM micrographs showing single folded SQB particles of the samples noted on the gel in *A*. Scale bars are 50 nm.

To determine whether there was any difference in the staple incorporation and folding quality with reused staple strands, we measured the presence of features on the origami. We initially examined a pair of 3′ ssDNA 16-nt handles on the extreme corners of the flat face of the SQB (i.e. the short light orange squiggles in Fig. 1Bi). We created two distinct cargoes with a complementary 3′ ssDNA antihandle, including a DNA origami barrel^14^ and a DNA nanocube^15^. As an example, we selected various purified SQB samples (depicted in Fig. 2D) for incubation: specifically, for the nanocubes, we chose SQB 0, 3, 7, and 11; whereas for the barrels, we opted for SQB 0, 6, and 11. We indirectly quantified the incorporation of the handle on the SQB by counting the number of cargoes conjugated to single particles in TEM images (Fig. 4A–B). Encouragingly, the relative conjugation of either the nanocube or barrel to the SQB was similar regardless of the number of times which the staple strands were reused and replenished (Fig. 4C). For instance, about ∼73% and ∼9% of the particles were bound with one and two nanocubes respectively, with this remaining nearly constant for the DNA origami folded using fresh staple strands versus those reusing the strands in three, seven, and eleven consecutive cycles.

**Figure 4.**
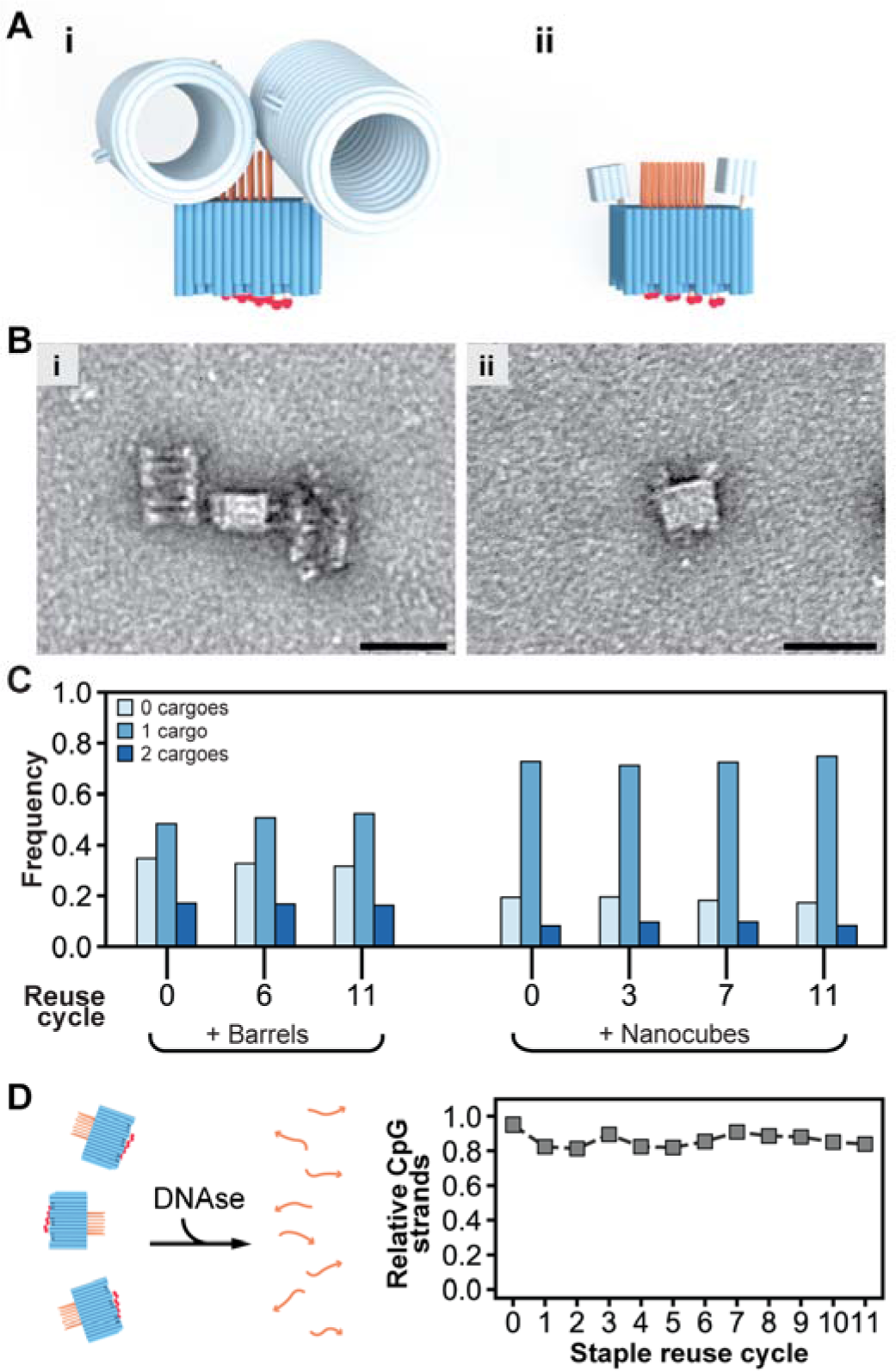
Programmed features on the staple-strand ends are successfully displayed on DNA origami therapeutics folded from reused strands with replenishment. **Ai–ii**, models of the barrel and nanocube cargoes attached to the SQB to assess incorporation efficiency of two handle strands. **B**, representative TEM micrographs of (**i**) barrels and (**ii**) nanocubes attached to the SQB. **C**, the relative frequency with which either zero, one, or two cargoes were observed in TEM micrographs of single SQB particles made from reused staple strands. N_particles_ = 570, 612, and 549 for SQB0, SQB6, and SQB11 respectively for the barrels. N_particles_ = 1100, 1288, 1821, and 1521 for SQB0, SQB3, SQ7, and SQ11 respectively for the nanocubes. **D**, the relative fraction of CpG strands remaining after DNase digestion of SQB samples made with reused staple strands.

We used the pairwise conjugation data in Fig. 4C to compute the probability that a cargo was attached to a single handle (see Supplementary Text 2). We determined there was ∼40% chance that a barrel would be bound to one of the handles tested on the SQB, irrespective of whether the origami was folded from fresh or reused staple strands. This computed probability might be representative of the incorporation efficiency of staple strands and their 3′ features on the origami, though there are several confounding factors that make it difficult to judge the true incorporation efficiency of all the single staple strands individually from solely this data, as explained in Supplementary Text 2. Nonetheless, the similarity in the relative conjugation frequencies for the SQB made using reused strands to its counterpart made with fresh staple strands suggests similar incorporation of the 3′ handle across all the samples tested. These observations suggest that origami folded using staple strands that were reused multiple times could be of sufficient quality for the validation of therapeutic DNA origami designs.

The other assay we did to determine if there was any difference in the staple incorporation between the samples in the various reuse rounds, was measuring the presence of CpG which is one of the most important features of the vaccine. The CpG strands are synthetic and this allows them to remain intact to the enzymatic activity of the DNase enzyme. This feature gives us the opportunity to perform a quantitative assay on the presence of this ODN on the SQB. We incubated the samples with DNase (see Method 11) and this enzyme digested all the origami leaving in solution just the strands of CpG (see Fig. S5). This was then quantified through gel densitometry giving as a result a stable rate of incorporation differing around 15% between the cycles (Fig. 4D).

To gain intuition of why reused strands could make well formed origami, we created a stochastic model to determine how staple excesses, impurities, and replenishment might influence the overall quality of the strand pool (Supplementary Text 3). We tested these experimental parameters in the model and qualitatively compared the results to the data in Fig. 2. From inspection of Fig. 2, we concluded that there is some amount of bias favoring the incorporation of full length staple strands during origami folding. Quantitative measurements of exactly how much the staple pool was degrading was beyond the scope of this manuscript. However, our stochastic model provides intuition of how strand quality might degrade through such cycling (Fig. S6A–B). Our stochastic model also illustrates how replenishment is critical to preserve the purity of the staple strands to yield origami with large proportion of full length staple strands over multiple cycles of reuse, and how larger excesses of staple strands when they are not replenished after each folding could slow the lessening of the quality of the origami over sustained reuse (Fig. S6C–D, leftward plots). Conversely, our model illustrates how replenishment is able to largely preserve the quality of the staple pool over multiple cycles of reuse regardless of whether the modeled folding is conducted with two-, five-, or ten-fold excesses of staple strands (Fig. S6C–D, rightward plots).

## CONCLUSION

This study showed that a DNA origami structure could be properly formed by folding it by reusing the excess staple oligonucleotides from prior folding experiments, so long as the staple strands were replenished with an equivalent amount of fresh staple strands to compensate for those lost during the previous folding experiment and the DNA ethanol purification process. We synthesized a proof-of-concept DNA origami therapeutic with satisfactory incorporation of cargoes by reusing the excess staple strands up to eleven times. A simplistic model of folding illustrates the extent to which the experimental process could bias the selection of full-length strands into the origami, which could explain why we observed high quality product formation despite the presumed gradual decrease in the purity of the staple pool. Reuse and replenishment of staple strands can significantly lessen the amount of staple strands that must be purchased to fold DNA origami. In turn, this could significantly lessen the costs of folding milligram amounts of DNA origami that would be needed to prototype and validate potential origami therapies in laboratories.

## AUTHORS CONTRIBUTION

Conceptualization, YCZ, CMW, GI; Methodology YCZ and GI; Validation, GI and CMW; Formal Analysis, CMW; Investigation, GI; Data curation, GI, CMW and PL; Writing - Original Draft CMW and GI; Imaging with TEM – GI and JD; Writing - Review and editing CMW, GI, OJY, YCZ and WMS; Visualization, CMW and GI; Supervision, YCZ and WMS; Project Administration, GI and YCZ; Funding Acquisition, YCZ and WMS.

## DATA AVAILABILITY

The data underlying this article are available in the article and in its online supplementary material. The Python code for the stochastic model is available from CMW. upon request. Folding sequences are available from previous citations.

## SUPPLEMENTARY DATA

Supplementary data is available online at NAR.

## ACKNOWLEDGMENTS

We thank Kathleen Mulligan for assistance with TEM imaging.

