## Supporting Information for "Reusing excess staple oligonucleotides for economical production of DNA origami"

**Table of contents**

**MAIN TEXT	1**

ABSTRACT	2

INTRODUCTION	3

RESULTS AND DISCUSSION	5

CONCLUSION	12

METHODS	13

AUTHORS INFORMATION	21

AUTHORS CONTRIBUTION	21

FUNDING SOURCES	21

ACKNOWLEDGMENTS	21

ABBREVIATIONS	22

**SUPPORTING INFORMATION	26**

**Supplementary Text 1: Cost discussion and projections about savings for staple reuse	28**

Supplementary Text 1.1 — Discussion about the niche of where reuse of excess staple strands is most economically compelling	28

Supplementary Text 1.2 — Generalized cost projections to make origami with reused staple strands	28

**Supplementary Text 2: Computing the probability of single handle incorporation from the two-handle conjugation experiments in Fig. 4A–C	30**

**Supplementary Text 3: Stochastic model of folding origami from reused staple strands	31**

Supplementary Text 3.1 — Purpose of the model and conclusions versus the experimental data	31

Supplementary Text 3.2 — Explanation of how the stochastic model works	32

Supplementary Text 3.3 — Shortcomings of the stochastic model	33

**SUPPLEMENTARY FIGURES	35**

**SUPPLEMENTARY TABLES	41**

### **Supplementary Text 1**: Cost discussion and projections about savings for staple reuse

#### Supplementary Text 1.1 *— Discussion about the niche of where reuse of excess staple strands is most economically compelling*

The DNA strands to fold DNA origami are typically acquired from two sources in academic research labs. Firstly, the long ssDNA scaffold is often the genome from the M13 bacteriophage, which is either sourced from commercial vendors or by M13 phage replication in *E. coli*. Although this can be a significant fraction of the total material cost, there is the potential for low-cost sourcing through growth of host bacteria in a fermenter^1–4^. For the purposes of this manuscript, we focus on reducing the material cost of the staple strands only.

The multiple staple oligonucleotides are typically purchased from commercial chemical DNA synthesis vendors. For experiments requiring nanomole or larger scale production of DNA origami, these can represent an especially onerous cost when there are expensive chemical modifications included, and furthermore when these strands are desired to be incorporated into origami at high occupancy, therefore demanding a large molar excess of these during folding. Here is a hypothetical example to illustrate the point: if a staple strand with an azide functionality is desired with folding at 10× molar excess over scaffold, then the cost from a vendor (e.g. IDT) could be $200 per nanomole origami. If the target origami were to require ten of these modifications per particle, then the material cost for just these azide-modified strands would be $2000 per nanomole origami particles^5^. To put this cost into context, a typical mouse experiment in our laboratory could require about 4 nanomoles of origami to treat 50 mice, therefore the cost for these modified oligonucleotides for such an origami could be on the order of $10k per study (although this cost could be considerably lower per unit mass for industrial scale production due to economies of scale), versus a cost on the order of $3k for the animals and animal housing. Thus we were highly motivated to study recycling of excess oligonucleotides here in order to reduce this cost.

One consideration of reusing staples to make intermediate scales of therapeutic DNA origami is that it requires additional experimental steps, where users must separate, reserve, and purify the excess staple strands when they fold more origami. However, downstream processing workflows already require separating excess staples before final use of the origami. For example, our protocol required PEG purification of the therapeutic origami, regardless of whether it was made using fresh or reused staples (see Method 4). The leftover excess strands could be collected and stored in batches in 4°C storage, and purified from the PEG supernatant by ethanol precipitation (see Method 5) weeks later at one’s convenience when the need to fold additional SQB origami arises.

#### Supplementary Text 1.2 *— Generalized cost projections to make origami with reused staple strands*

The method of staple reuse allows substantial cost savings for intermediate-scale fabrication of DNA origami using staple oligonucleotides from commercial vendors. To illustrate these savings for a more detailed example, we considered the following calculations. The **c**ost of staple strands (nt = nucleotide) for a fixed amount of origami **p**er **f**olding (cpf) as a function of the staple excess is

| $cpf(staple\_excess) = (total\_/origami*/mo{l staple}nt)mol\_origami *molarstaple\_excess$ | 1) |
| --- | --- |

The cost of staple strands for multiple batches of origami without versus with reuse as a function of the the number of cycles (“num_cycles”) in addition to the first one (i.e. cycles of reuse) and staple excess is

| $no\_reuse(num\_cycles, staple\_excess) =cpf(staple\_excess)*num\_cycles +cpf(staple\_excess)$ | 2) |
| --- | --- |
| $with\_reuse(num\_cycles, staple\_excess) = \frac{cpf(staple\_excess)*num\_cycles}{staple\_excess/recovery efficiency}+cpf(staple\_excess)$ | 3) |

And the cost difference in staple strands for folding multiple batches of origami with fresh staple strands versus with reuse with replenishment is

| $cost\_difference = no\_reuse(num\_cycle, staple\_excess) - with\_reuse(num\_cycles, staple\_excess)$ | 4) |
| --- | --- |
| $cost\_difference = cpf(staple\_excess)*num cycles*(1 - \frac{1}{staple\_excess/recovery efficiency})$ |  |

From equation 4), it is apparent that greater cost savings are driven by two factors. Firstly, the savings scale linearly with the number of cycles (i.e. number of fixed size batches of origami folded) of staple reuse/replenishment. Secondly, the relative cost savings are increased when larger excesses of staple oligonucleotides are used in the folding reaction (as per the last term in the above equation). It follows that with many cycles of staple reuse, the cost savings will approach the worth of the excess staple strands, as compared to an equivalent amount of origami made using entirely fresh staple strands.

The percentage cost savings versus the number of reuse cycles are shown in Fig. S1A. After ten cycles of reuse using two-, five-, and ten-fold excess of staple strands, there is a ~45%, ~73%, and ~82% reduction in cost of the staple strands consumed, respectively. We also considered the cost of the staple strands per unit of origami versus the total milligrams of origami produced (Fig. S1B). The latter plot assumes a unit price of $0.33 USD per base at 100 nmole yield (as from IDT Inc.), which was a typical price of the staple strands purchased for this work . Given this synthesis price, and with each folding batch consisting of 40 mL using 50 nM scaffold and a ten-fold excess of staple strands, the unit cost of the fresh staple strands for a single batch (i.e. ~11 mg of origami) would be ~$53 USD per milligram of origami. With ten cycles of reuse (~120 mg of origami total), this cost can be lessened to ~$10 USD per milligram of origami. It should also be noted that the percentage cost savings with the reuse of staple strands are independent of the price of DNA synthesis. So long as staple strand synthesis is a significant expenditure — which is of particularly large magnitude when chemically modified strands are needed in large excess — staple reuse is a useful approach to lowering the total price of DNA origami.

### **Supplementary Text 2**: Computing the probability of single handle incorporation from the two-handle conjugation experiments in Fig. 4A–C

We sought to determine changes in the relative probability of decoration of handles of SQB particles from successive cycles of staple-strand reuse. In experiments in Fig. 4A–C, we counted the number of origami barrels or nanocubes attached to a pair of 3′ ssDNA handles on single SQB particles in TEM images.

However, it is possible to compute the probability of incorporating single handles from paired handle data if we make two simplifying assumptions: (1) Each handle feature has the same probability of being added successfully; and (2) The binding event of the barrel or nanocube to one SQB handle occurs independently of another cargo binding the other SQB handle. These assumptions allowed us to treat the data from Fig. 4C as follows:

$$P_{data}(2 cargoes bind) = P_{computed}(1 cargo binds) * P_{computed}(1 cargo binds)$$

$$P_{data}(no cargoes bind) = P_{computed}(1 cargo does not bind) * P_{computed}(1 cargo does not bind)$$

Had the SQB had only one handle to bind a single cargo, it would follow:

$$P_{computed, SQB w/ 1 handle}(1 cargo binds) + P_{computed, SQB w/ 1 handle}(no binding) = 1$$

$$= P_{data}(2 cargoes bind)^{1/2} + P_{data}(no cargoes bind)^{1/2} = 1$$

We computed this from the data in Fig. 4C (blue columns) to see if it matched the expectations above and observed the following (yellow columns):

|  | **Data from** Fig. 4**C** | | **P_data_(2 bound cargoes)^(1/2) + P_data_(neither cargo bound)^(1/2)** | | **P_computed_(1 cargo binds)** |
| --- | --- | --- | --- | --- | --- |
| **Experiment** | **P_data_(2 bound cargoes)** | **P_data_(neither cargo bound)** | **Computed from data in** Fig. 4**C** | **Expected** | **Computed from data in** Fig. 4**C** |
| **Barrel SQB0** | 0.170 | 0.347 | 1.002 | 1 | **0.413** |
| **Barrel SQB6** | 0.167 | 0.327 | 0.980 | 1 | **0.408** |
| **Barrel SQB11** | 0.162 | 0.315 | 0.964 | 1 | **0.403** |
| **Nanocube SQB0** | 0.082 | 0.192 | 0.724 | 1 | 0.286 |
| **Nanocube SQB3** | 0.096 | 0.193 | 0.750 | 1 | 0.310 |
| **Nanocube SQB7** | 0.097 | 0.180 | 0.735 | 1 | 0.311 |
| **Nanocube SQB11** | 0.082 | 0.170 | 0.699 | 1 | 0.287 |

The data followed the expectation for independent binding with similar probability at each handle somewhat closely with the barrels. It suggests there might be ~0.4 probability that one of the handles is able to capture a barrel (see the bold numbers in the rightmost red column. Consequently, there might be ~0.4 probability that the 3′ end of one of these handles is functionally present on the SQB origami.

However, there was a mismatch from this expectation with the nanocubes. We rationalized these differences based on the sample preparation method for using transmission electron microscopy and experiment design. For instance, the orientation in conjugated particles as settled on the flat carbon imaging substrate could have obscured the nanocube cargo under the larger footprint of the SQB. Consequently, we would have been unable to count the true number of times that the SQB was bound to either one or two of its nanocubes. Moreover, there might have been steric hindrance of handles on the SQB, where an initially bound cargo lowered conjugation efficiency to the remaining unbound handle. It is also conceivable that some amount of the cargoes became separated from the SQB during sample preparation techniques. Alternatively, the simplifying assumptions noted above might not actually match the true behavior of when cargoes bind to the SQB.

Further experimentation with different techniques would be required to validate the probability of successful staple and handle incorporation of single handles.

### **Supplementary Text 3**: Stochastic model of folding origami from reused staple strands

#### Supplementary Text 3.1 — Purpose of the model and conclusions versus the experimental data

To rationalize why reused strands can make well-formed origami, we created a stochastic model to determine how staple excesses, impurities, and replenishment might influence folding performance. In the model, a given staple strand in the total population of strands is in one of two states. First, its desired full-length; or second, flawed with some truncation to its length. The state of a particular strand determines the probability that it will be incorporated into an origami during folding, and we presumed there would be some bias favoring incorporation of full-length staple strands into the origami over truncated strands. We introduced a bias factor σ into the model and tested different degrees of selection for the full length stands, as shown in Fig. S6A.

Just for illustration purposes, we make the naive assumption (without justification) that the probability of proper folding is linearly related to the mean fraction of full-length staple strands incorporated per origami. To determine the bias σ that best resembled experimental observations given the naive assumption above, we ran the model using a two-fold excess of staple strands (Fig. S6B). Relatively high proportions of full-length staple strands were added to the origami over multiple cycles of reuse with no replenishment and little bias (i.e. σ = 1.000001), contradicting the data in Fig. 2Ai. We progressively incremented the bias σ to e, 10, 20, and 1×10^6^ and observed that the origami had decreasing proportions of full length staple strands with larger values of σ, as the staple strands were reused without replenishment (top row Fig. S6B). This decrease in full length strands could largely be mitigated by replenishing the strands, though more σ bias led to a gradual decline in the amount of full length strands with continued reuse (bottom row Fig. S6B).

In particular, the behavior of the model when σ was equal to 20 qualitatively resembled the data in Fig. 2A–B, motivating us to test the model using this σ bias and different excesses of strands. Larger excesses of staple strands in either five- or ten-fold excess with reuse and no replenishment were able to significantly lessen the decrease in full-length strands in the modeled origami, compared to when only a two-fold excess was used (leftward Fig. S6C). With replenishment, the proportions of full length strands in the origami were increasingly stable as higher excesses of strands were used (rightward Fig. S6C). Similarly, the standard excesses of strands (where some subset of the 277 strands were in either five-, ten-, or twenty-fold excess) exhibited similar behavior with and without reuse across multiple cycles (Fig. S6D). Taken together, the model suggests that larger excesses of strands in conjunction with replenishment could maintain the quality of reused staple strands to fold satisfactory origami, despite how the folding process preferentially extracts full length strands each folding cycle. ***Note: the reason we cannot make a quantitative determination about σ from the experimental data in this paper is explained further in*** Supplementary Text 3***.3.***

#### Supplementary Text 3.2 — Explanation of how the stochastic model works

We sought to rationalize our experimental results showing that reused staple oligonucleotides with replenishment could be used to fold DNA origami of high quality. We used a stochastic model to test how the relative purity of staple oligonucleotides might influence the probability that a given origami particle incorporates a full length staple. The model was coded in Python and simplified the DNA folding process as follows.

1) For the purposes of the model as tested within this manuscript, we consider the SQB origami design that is composed of a single scaffold strand and 277 distinct staple oligonucleotides that range in length from 32 to 58 nucleotides.
2) The model is initialized with some number of each staple oligonucleotide (e.g. 20,000 of each distinct strand) and with a number of scaffold strand molecules (e.g. 10,000). The number of scaffold molecules is selected so as to maintain some user-defined excess with respect to the staple strands (e.g. 2× as considered here).
3) The initial purity of the staple oligonucleotides is 0.994^(bp length - 1), which was conservatively estimated from the synthesis yield of oligonucleotides as described by the manufacturer Integrated DNA Technologies (IDT) on their website^6^. Purity is defined as the relative fraction of the staple strands which are their full designed length.
4) The folding process entails each scaffold strand extracting one of each of the 277 distinct staple oligonucleotides from the total pool of staple oligonucleotides. The probability that a scaffold strand will remove a full length ‘pure’ strand versus a truncated ‘impure’ strand is determined as some function of the purity of that particular strand in the staple oligonucleotide pool.

In particular, we tested the following model for the incorporation frequency of full length strands, using different relative bias factors favoring the full length staple strands. As σ → 1, the bias for full length strands over their truncated counterparts diminishes. And as σ → ∞, the bias for full length strands over their truncated counterparts becomes increasingly stronger.

a) $P(full length strand addition) = log_{\sigma}[(\sigma-1) \times purity + 1]$
b) $P(truncated strand addition) = 1 - P(full length strand addition)$

5) Remaining staple oligonucleotides that are not incorporated into DNA origami are reused repeatedly to fold additional DNA origami for some arbitrary number of cycles of reuse.
   a) With no replenishment, the number of scaffold strand molecules added is lessened accordingly to maintain the user-defined staple oligonucleotide excess.
   b) With replenishment, additional staple strands are added to replace the staple strands which were removed by origami in the previous folding step. The number of each staple oligonucleotide added is the difference between the number of each staple added versus the scaffold strand that was initially added (see step 2). The relative purity of the added replenishing strands is as described in step 3.
6) The relative proportion of full length ‘pure’ staple oligonucleotides added to each origami versus the extent of reuse was measured and used as an indicator of the possible quality of the folded structures.

#### Supplementary Text 3.3 — Shortcomings of the stochastic model

We note that our model is an extreme simplification of the origami folding process. It lacks granularity about multiple details of the folding pathway and strands themselves. These limitations include the following.

1) The data computed in the model versus the gel densitometry data are measurements of two different phenomena — that while likely correlated and informative of one another — are not a like-for-like comparison. The model data in Fig. S6B–D are the relative number of full length strands incorporated into the *in silico* origami. Conversely, the gel densitometry data in Fig. 2 assesses the relative proportion of DNA origami that migrate at the expected molecular weight for the SQB particle.

The observation of slower gel migration of the origami when the staple strands are reused without replenishment in Fig. 2Ai is conceivably due to an accumulation of truncated strands in the SQB product. Nonetheless, full quantitative connection of the model to experimental data would require collection of more granular experimental data. For instance, in another study the experimenter could perform next-generation sequencing of the folded origami particles to determine the proportion of full versus truncated staple strands and their relative location in the folded structures.

2) There are no sequence information considerations of the staple oligonucleotides or design details for a particular origami shape. It is conceivable that there could be some sequence or location dependence on thermodynamics of strand binding for a particular strand to be successfully incorporated.
3) The model did not consider any flaws to the strands beyond truncations to their one dimensional length. It is likely that there are other types of flaws that might be present in a set of oligonucleotides synthesized using phosphoramidite chemistry.
4) Individual failure modes of staple strand incorporation is not considered. For instance, a full length strand could be misincorporated such that the 3´ or 5´ end feature is not displayed on the structure. Moreover, a given strand (either full length or truncated) in a particular origami particle might also not be added at all.
5) Cooperativity between individual staple strands during folding is not considered. For instance, the binding of certain staple strands in some other location within the origami could influence the probability that a given staple at another proximal location is subsequently incorporated.

Nonetheless, our model superficially followed the experimental behavior of folding with reuse and replenishment. It gave a sense of the bias with which DNA origami folding might favor selection of full length strands over truncated flawed impurities. Moreover, the model suggested that using larger excesses of staple strands could make successful origami folding more robust to multiple cycles of reuse and replenishment.

### SUPPLEMENTARY FIGURES

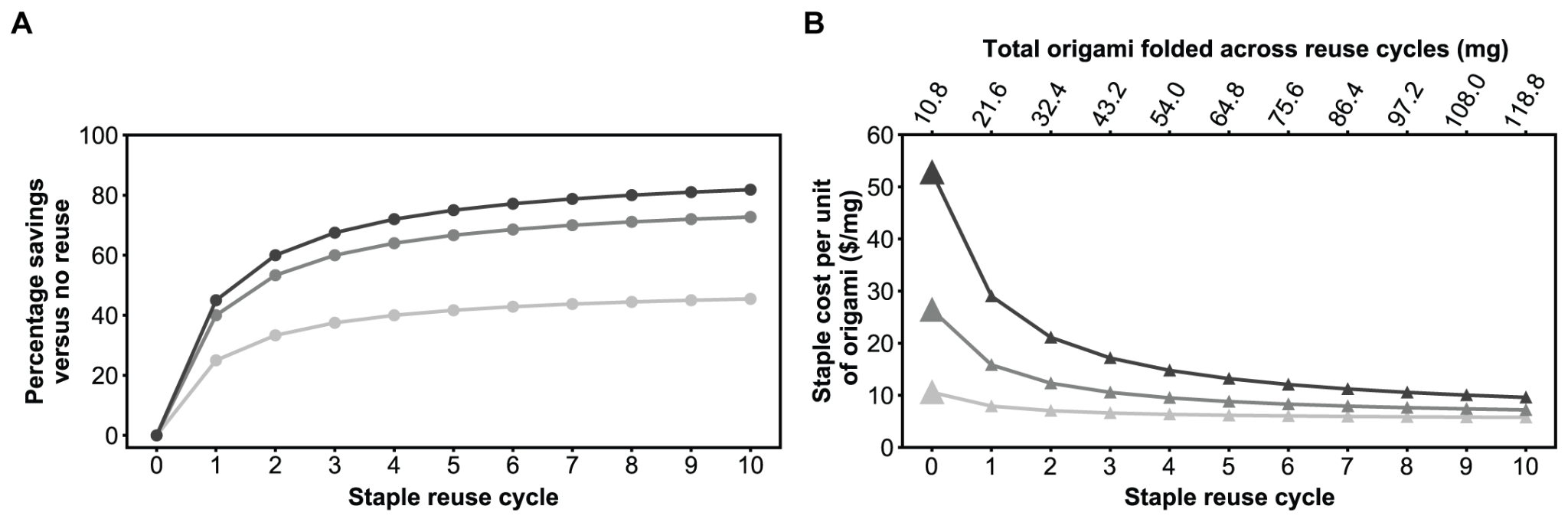

**Figure S1** Cost savings of reusing excess staple strands to fold DNA origami. The black, medium gray, and light gray lines correspond to 10×, 5×, and 2× excesses of staple strands with respect to the scaffold in the folding reaction. **A**, Percentage savings in synthetic staple strands over multiple cycles of reusing and replenishing excess staple strands versus folding an equivalent amount of origami using only freshly purchased synthetic staple strands. Greater cost savings are realized with **B**, the US dollar cost of synthetic staple strands per milligram of origami versus the total amount of DNA origami folded. By comparison, the cost of staple strands per unit of origami remains fixed if only fresh staple strands are used (i.e., at the rate indicated with the enlarged data points). The cost projection assumes 8634 nucleotides of oligos at a 100 nmole scale at $0.33 USD/base, which is the price as listed by IDT at the time of this writing. Each x-axis increment of 10.8 mg represents the folding of 40 mL of origami using a p8634 scaffold at 50 nM concentration.

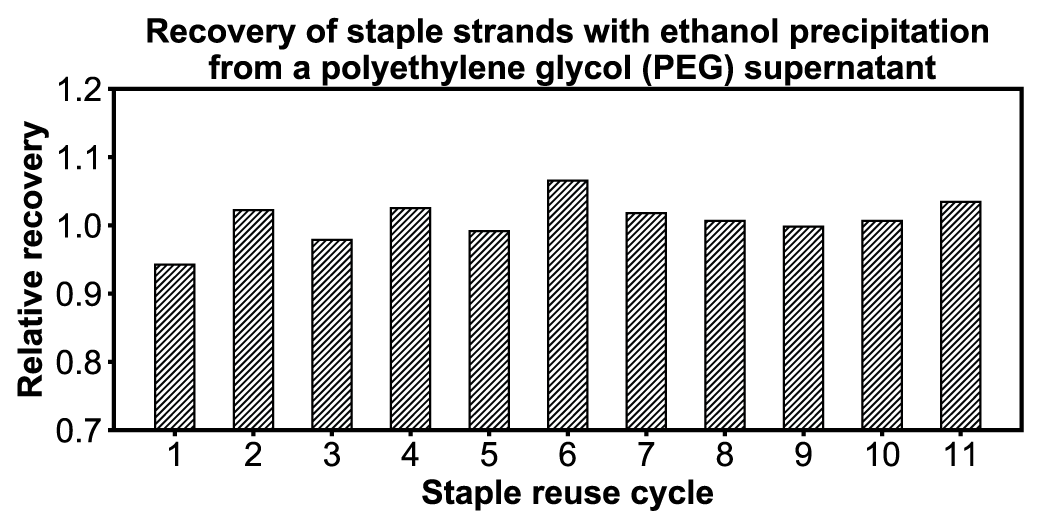

**Figure S2**. The relative recovery of excess staple strands from the leftover PEG supernatant using ethanol precipitation so that the staple strands may be reused to fold more DNA origami. The DNA concentration of the staple strands in the input PEG buffer and the output buffer once the strands were dissolved back into a buffer was measured using a nanodrop spectrophotometer. There were comparable amounts of strands in the input material and output product which indicated that ethanol precipitation could recover strands from a PEG supernatant with little loss of material.

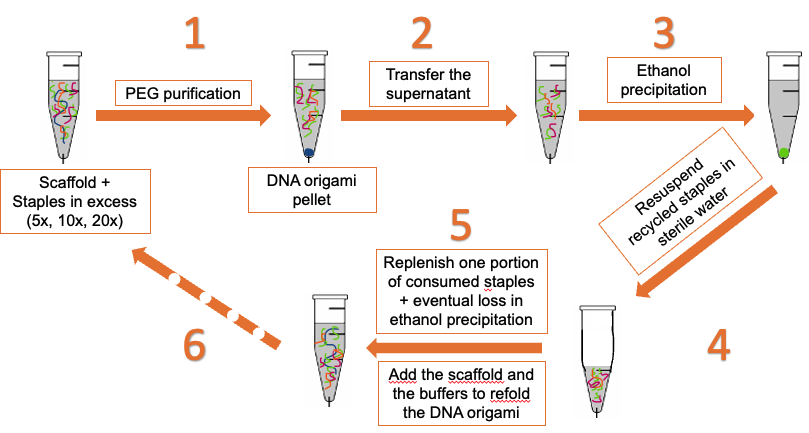

**Figure S3** Flowchart of methodology to fold, recover, replenish, and reuse excess staple strands oligonucleotides. **1.** The folding solution after the thermal ramp cycle needs to be PEG purified in order to get rid of all of the staple strands excess. This step will allow the separation of the components of the solution, at the bottom of the tube there will be the pellet containing the folded molecules and left in solution there will be the excess of staple strands. **2.** The supernatant will be transferred to another tube and in the original one the pellet will be resuspended with the appropriate buffer. **3.** The solution in the new tube will face ethanol precipitation, method used to precipitate all the staple strands at the bottom of the tube. At this point all the staple strands will be precipitated in a pellet, the ethanol and the salt in solution need to be washed off. **4.** Once done one so the staple strands will be brought to solution with sterile water at the approximate concentration desired. The staple strands are now ready to be reused. **5.** Then it’s necessary to add the consumed staple strands for the previous folding and the eventual loss in the steps due to ethanol precipitation or manual errors. The other components necessary for the folding will be added and **6.** the solution will go under the thermal ramp to face another round of folding.

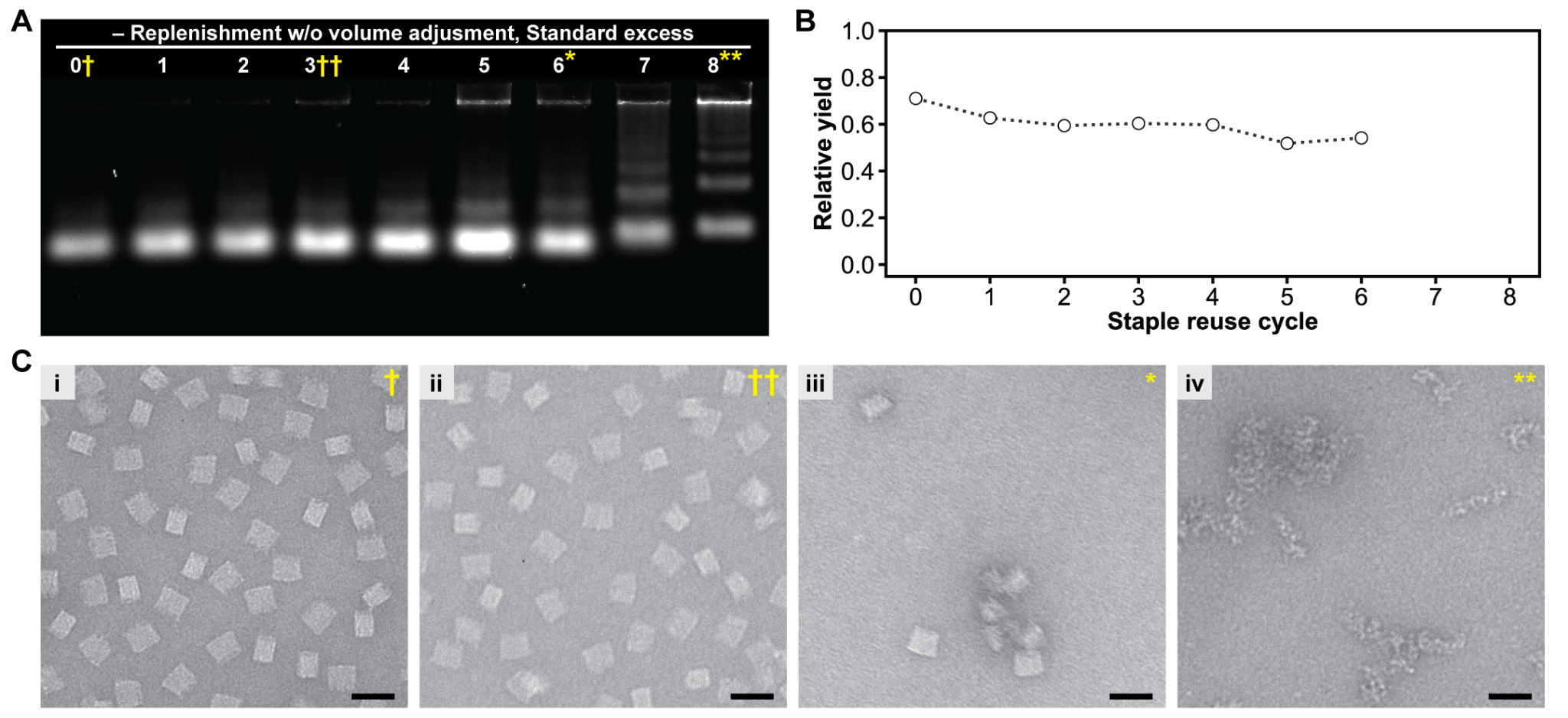

**Figure S4** SQB folding results with reuse of excess staple strands, without replenishment of excess staple pools and adjustment to folding reaction volume in the subsequent reuse cycle. **A**, agarose gel showing that there is a substantial slowdown in the migration of the SQB after six or more rounds of reuse without replenishment and volume adjustment. The number of reuse cycles is denoted with the numbers above each lane of the gel. **B**, the relative yield of the SBQ was calculated from *A* with agarose gel densitometry except for lanes 7 and 8 due to TEM evidences. **C**, TEM micrographs of the SQB with zero (***i***), three (***ii***), six (***iii***), and eight (***iv***) cycles of reuse with no replenishment and adjustment to reaction volume. This lessening of quality of the SQB origami is attributed to poor folding of the origami due to depletion of select staple strands from the reused strands and/or incorporation of truncated staple strands of poor quality. By contrast, the SQB origami could be satisfactorily folded from reused staple strands when each folding reaction was replenished with an equivalent of staple strands that was depleted in the prior folding, as shown in Fig. 2D–E. Staple strand excesses were the standard amounts, as noted in the text with regard to their therapeutic importance. Scale bars in ***C*** are 50 nm.

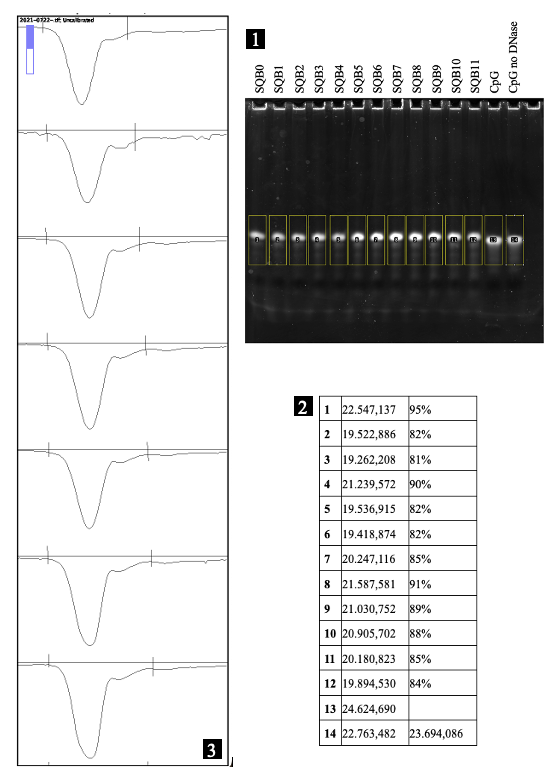

**Figure S5** For estimating CpG conjugation efficiency, we utilized the ImageJ program. We delineated each lane of a PAGE gel using a rectangular shape, as demonstrated in Fig. S5.1. We analyzed each band of interest from the experimental samples, which were treated with replenishing, and compared them with the last two lanes containing the controls: CpG digested with DNase and untreated CpG. After tracing the boxes on the gel with consistent dimensions to ensure accurate measurements, we quantified the intensity of each band.ImageJ provides a feature to display intensity curves, allowing the operator to select the appropriate area with two lines delimiting the perimeter (Fig. S5.3). Using the "wand tracing tool," we measured the areas, and the corresponding numbers were generated and recorded in the second column of the table in Fig. S5.2. The last two lanes served as our controls; we averaged their values and utilized that average (located in the bottom right corner of the table) for further calculations. We individually divided the intensity of each lane by the averaged value to derive the conjugation efficiency rate, expressed as a percentage in the third column. The average conjugation efficiency rate was then calculated, resulting in an efficiency of 86%.

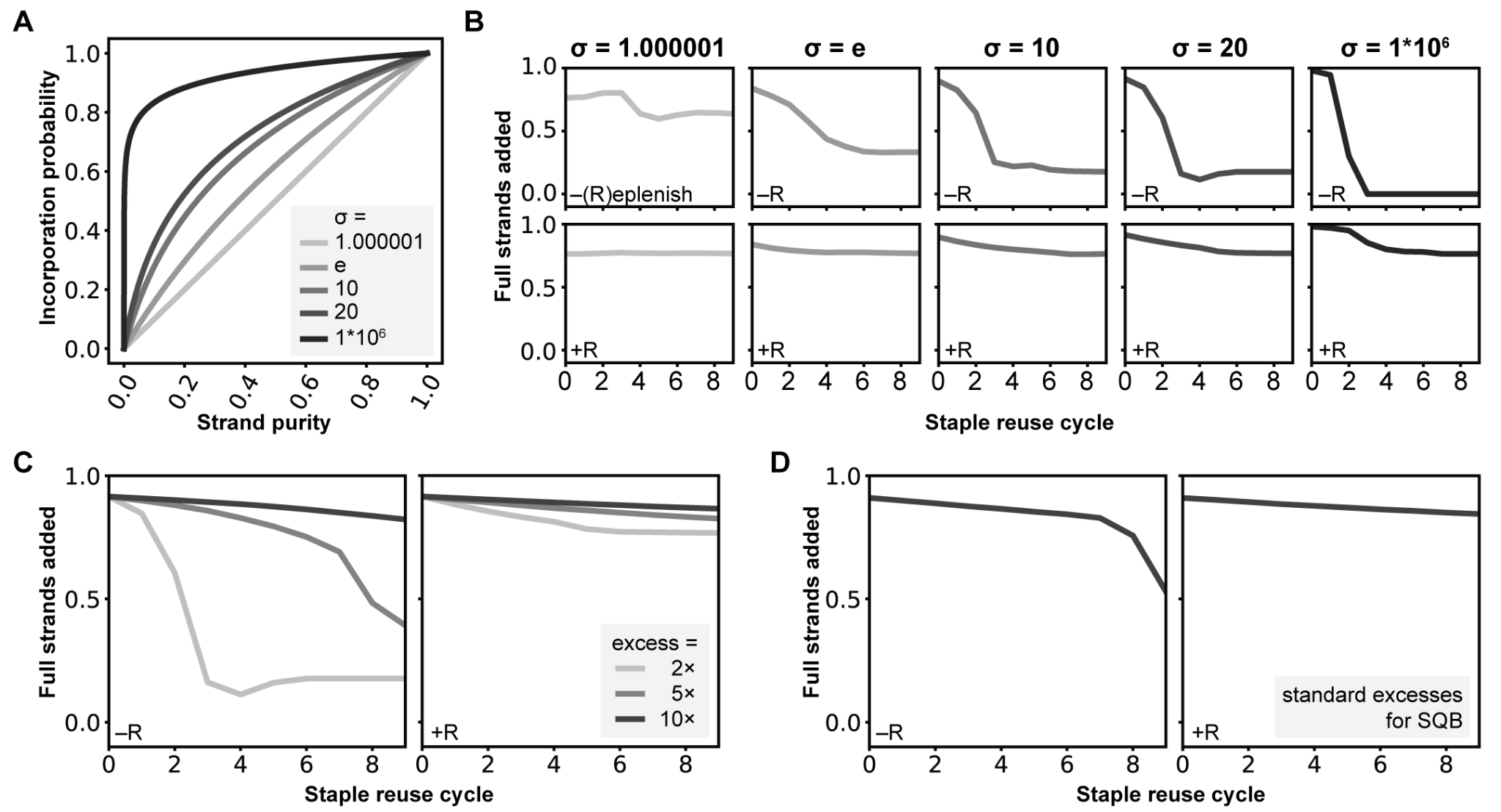

**Figure S6** Stochastic model of staple addition into DNA origami when reusing of excess staple strands with and without replenishment. **A**, several plausible assumptions for the bias σ that full-length staple strands are added to the origami versus purity of the staple pool. **B**, staple incorporation into the origami with a 2× excess of strands using different amounts of bias. **C**, staple incorporation using either 2×, 5×, or 10× excess of strands with one bias value (σ = 20). There is a precipitous drop in the proportion of full-length strands added after two rounds of reuse with no replenishment, using a 2× excess of strands. **D**, staple incorporation into the origami using the standard excesses of strands as determined by their therapeutic relevance in the SQB with one bias value (σ = 20). Labels **–R** or **+R** in *B*, *C*, and *D* indicate without and with replenishment, respectively.

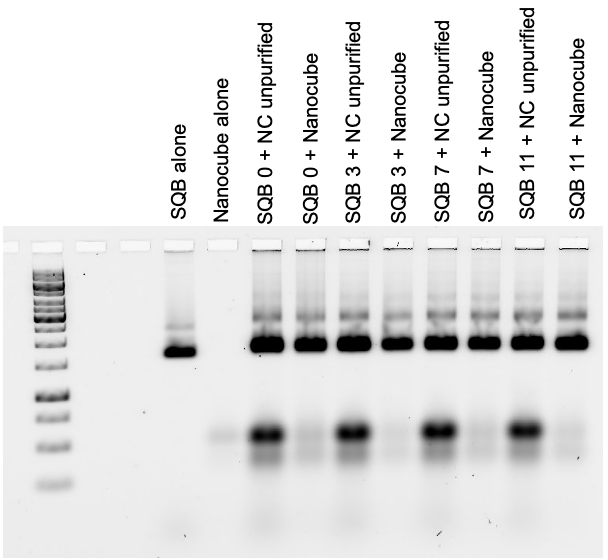

**Figure S7** Agarose gel electrophoresis of nanocubes (NC) conjugated with the SQB sample, as depicted in Figure 3A, before and after purification.

### SUPPLEMENTARY TABLES

**Table S1** Standard staple strand excesses for folding the SQB.

| **Staple type** | **Staple name** | **Excess vs. scaffold** | **[each staple] in 2× staple stock (μM)** | **[each staple] in final reaction (μM)** |
| --- | --- | --- | --- | --- |
| Core staple strands | A, B, C | 5× | 1.0 | 0.5 |
| Handles | Juno, Jupiter | 10× | 2.0 | 1.0 |
| Additional handles | Diana | 10× | 2.0 | 1.0 |
| Functional staple strands | CpG | 20× | 4.0 | 2.0 |

**Table S2** Our negotiated purchase prices per μl of the various types of staple oligonucleotides necessary to fold the SQB origami.

| **Staple type** | **Prices ($)** | **Volume (μl)** | **$ per μL** |
| --- | --- | --- | --- |
| Plate strands A | 2500 | 12600 | 0.20 |
| Plate strands B | 2500 | 15200 | 0.16 |
| Plate strands C | 2500 | 11200 | 0.22 |
| Plate with handles | 2000 | 6400 | 0.31 |
| Plate CpG | 4500 | 3600 | 1.25 |

**Table S3** Costs of a regular 40ml folding for animal studies differentiated per types of staple strands

| **Staple type** | **μl per 40 mL batch** | **$ per μl** | **$ per 40 mL batch** |
| --- | --- | --- | --- |
| Plate strands A | 2520 | 0.20 | 500 |
| Plate strands B | 3040 | 0.16 | 500 |
| Plate strands C | 2560 | 0.22 | 571.4 |
| Plate with handles | 2240 | 0.31 | 700 |
| Plate CpG | 2880 | 1.25 | 3600 |
| **Total** | 13240 | \ | 5871.43 |

**Table S4** Costs to replenish one part of the excess for resuing the staple strands in a 40 mL folding

| **Plates types (excess)** | **40ml folding** | **1 part** | **$** |
| --- | --- | --- | --- |
| Plate strands A (5×) | 2520 | 504 | 100 |
| Plate strands B (5×) | 3040 | 608 | 100 |
| Plate strands C (5×) | 2560 | 512 | 114.3 |
| Plate with handles (10×) | 2240 | 224 | 70 |
| Plate CpG (20×) | 2880 | 144 | 180 |
| **Total** | 13240 | \ | 564.3 |

**Table S5** Materials required to perform ethanol precipitation

|  | **Ethanol** | **Sodium Acetate 3M** |
| --- | --- | --- |
| **Price per bottle ($)** | 4 | 80.20 |
| **Volume per bottle (ml)** | 1000 | 100 |
| **Volume required for procedures (ml)** | 31.5 | 4 |
| **Price for materials used ($)** | 0.126 | 5.00 |
| **Total staple strands + materials** | \ | 569.4 |

**Table S6** Standard staple strand excesses for folding the Barrel

| **Staple type** | **Staple name** | **Excess vs. scaffold** | **[each staple] in 2× staple stock (μM)** | **[each staple] in folding reaction (μM)** |
| --- | --- | --- | --- | --- |
| Inner middle staple strands | PS-1 | 5× | 1.0 | 0.5 |
| Jupiter handle | PS -2A (handles for Cy5) | 5× | 1.0 | 0.5 |
| Juno handle | PS - 2B (handles for CpG) | 20× | 4.0 | 2.0 |
| Neptune handle | PS - 2C (handles for dsRNA) | 20× | 4.0 | 2.0 |
| Inner miniscaffold strands | PS - 3 | 5× | 1.0 | 0.5 |
| Outer H172 miniscaffold strands | PS-4 | 20× | 4.0 | 2.0 |
| Inner middle staple strands | PS - 5 | 5× | 1.0 | 0.5 |
| Ori handle or pool handle (not sure) | PS - 6 | 20× | 4.0 | 2.0 |
| Fluorophore | Cy5 | 5× | – | 2.5 |

**Table S7** Conjugation rates for SQB with nanocubes

| **SQB n°** | **0 Nanocube** | **1 Nanocube** | **2 Nanocubes** |
| --- | --- | --- | --- |
| **SQB 0** | 19.2% | 72.6% | 8.2% |
| **SQB 3** | 19.3% | 71% | 9.6% |
| **SQB 7** | 18% | 72.3% | 9.7% |
| **SQB 11** | 17% | 74.8% | 8.2% |

**Table S8** Conjugation rates for SQB with barrels

| **SQB n°** | **0 Barrel** | **1 Barrel** | **2 Barrels** |
| --- | --- | --- | --- |
| **SQB 0** | 34.7% | 48.2% | 17% |
| **SQB 6** | 32.7% | 50.7% | 16.7% |
| **SQB 11** | 31.5% | 52.3% | 16.2% |

**REFERENCES – SUPPORTING INFORMATION**

6.	Oligonucleotide synthesis: Coupling efficiency and quality control | IDT. *Integrated DNA Technologies* https://www.idtdna.com/pages/education/decoded/article/oligo-synthesis-why-idt-leads-the-oligo-industry.
